# Mass photometry reveals stoichiometry and binding dynamics of bispecific tetravalent anti-VEGF-PD-1 antibody ivonescimab

**DOI:** 10.1101/2025.09.05.674400

**Authors:** Nina Jajcanin Jozic, Josh Bishop, Alexander Ross O’Shea, Gogulan Karunanithy, Catherine Lichten, Svea Cheeseman

## Abstract

**Background:** The bispecific antibody ivonescimab targets programmed cell death protein 1 (PD-1) and vascular endothelial growth factor (VEGF). Recent clinical trials have shown it has greater efficacy against PD-L1 positive non-small cell lung cancer than pembrolizumab (Keytruda), a frequently prescribed anti-PD-1 monoclonal antibody. Ivonescimab binds to two VEGF and two PD-1 molecules, with complex formation through higher-order structure formation (or ‘daisy chain’ binding). However, the binding stoichiometries and interaction dynamics of ivonescimab with VEGF and PD-1 have not been characterized in depth.

**Methods:** We used mass photometry (MP) and kinetic modelling to analyze these interactions, quantifying the complexes formed and their affinities. Dissociation constants (K_D_) for ivonescimab’s binding to VEGF and PD-1 were calculated from equilibrium counts and real-time measurements, respectively.

**Results:** VEGF drove oligomerization of ivonescimab, which bound VEGF predominantly in a 2:2 stoichiometry, with K_D_=0.08 nM. Higher-order oligomeric complexes, present only at low abundance, displayed markedly weaker affinities (3.17 nM; 1.29 nM). Ternary complexes of ivonescimab with its two targets consistently presented two PD-1 antigens for each ivonescimab molecule, with a 1.66 nM K_D_ for the binding of the first PD-1 and the slightly stronger 0.89 nM for the second PD-1 molecule.

**Conclusions:** MP confirmed VEGF-induced ivonescimab oligomerization and revealed that dimers, not higher-order structures, were the most stable stoichiometry. MP enables detailed analysis of antibody–antigen interactions, even for bispecific antibodies that interact with antigens with complex stoichiometries.

**Statement of significance:** We report the first application of mass photometry to characterize the binding of the clinically promising bispecific antibody ivonescimab to its targets, VEGF and PD-1. The most stable assembly is a dimeric complex, not higher-order, as expected – advancing understanding of ivonescimab and demonstrating mass photometry’s value for complex biologics analysis.

## Introduction

Ivonescimab is a first-in-class bispecific antibody developed by Akeso Biopharma and Summit therapeutics. It is considered an important therapeutic innovation^1^, as it binds simultaneously to two critical cancer targets: *Programmed death protein-1* (PD-1) and the *vascular endothelial growth factor* (VEGF).

PD-1 and its ligand PD-L1 are major immune checkpoint proteins that downregulate the immune response when active, enhancing tumor cell survival. Immune checkpoint inhibitors that target the PD-1/PD-L1 axis (e.g. pembrolizumab (Keytruda) for PD-1 and atezolizumab (Tecentriq) for PD-L1) are important cancer treatments and are known to improve patient outcomes^2^. However, many patients develop resistance to these drugs, resulting in tumor recurrence^3,4^.

VEGF, meanwhile, is a protein that plays a crucial role in angiogenesis. In tumorigenic processes, VEGF is frequently upregulated within the tumor microenvironment (TME), which supports the formation of new blood vessels that provide cancer cells with oxygen and nutrients. For this reason, VEGF is another frequent target of anti-cancer drugs, but it is also associated with acquired resistance in patients^5^.

To avoid acquired resistance, a frequent treatment strategy combines anti-VEGF drugs with immune checkpoint inhibitors^6^. Conventional combination therapies require administering separate anti-PD-1/PD-L1 and anti-VEGF antibodies. Ivonescimab represents an innovation because it combines bevacizumab (anti-VEGF) and penpulimab (anti PD-1) in the same molecule^7^. Moreover, ivonescimab is engineered so that its two domains act in a cooperative manner, where binding at one domain increases the other’s binding affinity^7,8^. Its proposed method of action is that the bispecific antibody first binds to VEGF, which is abundant in the TME, leading to the formation of higher-order structures of multiple antibodies and antigens^7^ (‘’daisy chaining’). It is thought that these higher-order structures then bind to PD-1 on the T-cell surface with higher affinity (mainly as a result of a lower dissociation rate constant, k_off_, as compared to PD-1 antibodies alone)^7^.

Ivonescimab has thus far demonstrated promising therapeutic potential. Its first trial in humans showed efficacy signals and acceptable safety profiles^9^. HARMONi-2, a randomized, double-blind, phase 3 trial across 55 hospitals in China, showed that ivonescimab significantly improved progression-free survival (PFS) compared with pembrolizumab (Keytruda) in previously untreated patients with advanced PD-L1 positive non-small cell lung cancer^10^.

Despite these promising clinical results, ivonescimab has yet to be characterized biochemically in depth, and crucial details about its binding mechanics, oligomerization behavior and typical binding stoichiometries with both targets are still missing. A key reason for this knowledge gap is the lack of well-established analytical techniques that can identify multiple antibody-antigen complexes with varying stoichiometries and quantify their binding affinities. Conventional techniques such as biolayer interferometry (BLI) and surface plasmon resonance (SPR) are ensemble measurements and optimized for analyzing 1:1 interactions. They are less suited to analyzing binding in multi-specific antibodies.

To address this gap, we applied mass photometry (MP) to characterize the interactions of ivonescimab with its two targets. MP is an analytical technique that measures the interference of the light reflected by a measuring surface illuminated by a laser and the light scattered by particles in contact with that surface. The resulting signal or “contrast” is proportional to the molecular mass of each measured particle^11^. MP reports the mass distribution of a sample, providing an overview of all the components present and their complexes, with single-molecule resolution. MP results have been shown to align with results from size-exclusion chromatography (SEC) as well as analytical ultracentrifugation (AUC)^12^. Using this technique, we characterized the binding dynamics and stoichiometry of the interactions between ivonescimab and its two targets. Our results provide an overview of the interactions between ivonescimab, PD-1 and VEGF, revealing a range of different complexes with varying stoichiometries and suggesting that a 2:2:4 ternary complex between the antibody and its two targets is the predominant assembly state.

## Materials and methods

### Materials and Reagents

Ivonescimab was purchased from Cambridge Bioscience (cat no HY-P99675), Human PD-1 / PDCD1 Protein, His Tag was purchased from AcroBioSystems (PD1-H522a). Recombinant human VEGF 165A protein (Active) was purchased from Abcam (ab259412). Penpulimab was purchased from Cambridge Bioscience (cat no HY-P99108). Pembrolizumab (Keytruda) was purchased from MedChemExpress (HY-P9902), MassFerence^TM^ P1 calibrant was obtained from Refeyn Ltd., and Dulbecco’s phosphate-buffered saline (DPBS) was from ThermoFisher Scientific (14190144).

### Affinity and kinetic analysis of VEGF and PD-1 binding to ivonescimab

All measurements were done on a TwoMP mass photometer (Refeyn Ltd.), using MassGlass^TM^ UC slides. The slides were additionally cleaned, as recommended by the manufacturer for imaging samples containing molecules with mass <50 kDa. Briefly, the slides were cleaned by dipping them in 100% ispropanol followed by rinsing in mqH_2_O. This step was repeated 3-4 times, after which slides were air gun dried. All the reactions were performed at room temperature (RT), and samples were diluted in PBS prior to measurement, where indicated in the Results section. Also, prior to each set of measurements, a mass calibration was performed using the MassFerence™ P1 calibrant (Refeyn Ltd.). All data were acquired using AcquireMP v2025 R1 and processed using DiscoverMP v2025 R1 software.

For K_D_ analysis of the ivonescimab and VEGF complexes, both were mixed at 50 nM for the 1:1 molar ratio, and at 50 nM and 100 nM, respectively for the 1:2 molar ratio. Here and throughout the manuscript, concentrations refer to the monomer concentration. The prepared reactions were incubated at RT for the period indicated, after which quick dilution was done directly in a PBS droplet and the sample was immediately imaged (the dilution factor depends on the initial concentration in the mixture and was chosen to optimize the measurement in accordance with guidance in the instrument’s manual.)^13^ For each experiment, population masses and relative abundance were obtained from automated Gaussian fitting of the mass distribution in the DiscoverMP software. For calculating the K_D_ from MP counts, as described in Soltermann et al.,^14^ the concentrations used were those present in the droplet. Dissociation data (not shown) confirmed that higher-order complexes had not dissociated within 30 min after dilution to nanomolar concentration that is around or higher then estimates KD, confirming the system was in equilibrium. The K_D_ calculations done here are described in detail elsewhere^12,13^. In short, the measured counts were converted into molar concentrations, using a conversion factor, *f_conversion_*that was calculated for ivonescimab using the following equation:

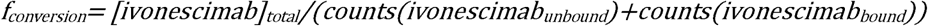

where counts(ivonescimab_unbound_) and counts(ivonescimab_bound_) are the counts obtained from the Gaussian fits. When counting total ivonescimab molecules, we corrected the counts by taking into consideration the stoichiometry of each complex. As the VEGF mass is close to the detection limit of the instrument, the concentration was calculated using mass balance rather than using a conversion factor, following this calculation:

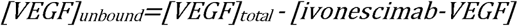

The K_D_ was then obtained from the standard equilibrium equation:

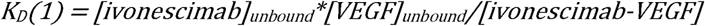

For other reactions and in the cases of multivalent binding, the same principle for calculating K_D_ was applied.

For the kinetic analysis of ivonescimab-PD-1 complex formation, a 1:4 mixture of ivonescimab and PD-1, with 5 nM ivonescimab, was prepared. The mixture was incubated for the indicated periods (between 10 sec and 20 min, then measured (with or without further dilution, as indicated in the Results). Data were recorded for 60 s, with a regular field of view setting in AcquireMP, with a frame rate of _≈_ 500 Hz, and frame binning of 10. For each experiment, a Gaussian Mixture Model consisting of three Gaussians centered to known masses (210, 240, 270) with a single shared sigma was automatically fitted to the mass histogram. Event counts for PD-1, ivonescimab, ivonescimab-PD-1, and ivonescimab-(PD-1)_2_ were extracted by integrating the individual Gaussian fit components. Higher-order assemblies (involving more than one ivonescimab molecule) were excluded from the analysis due to their low abundance. As for the interaction with VEGF, as unbound PD-1 is near the instrument’s detection limit, its concentration was calculated using mass balance, following this calculation:

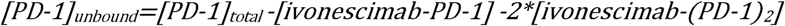

The midpoint of each acquisition was used as the time coordinate for kinetic modelling. The binding reaction between ivonescimab and PD-1 was modeled as a two-step process involving the sequential addition of PD-1 monomers:

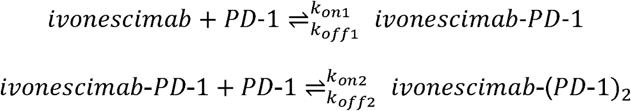

The kinetic model was defined by the following system of ordinary differential equations:

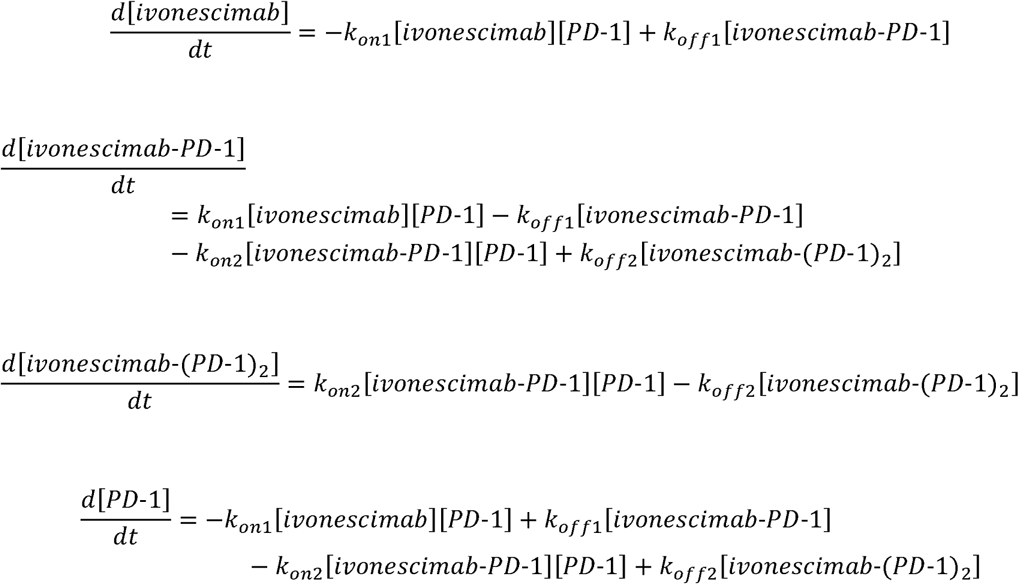

Kinetic rate constants were obtained by performing a least-squares optimization between the simulated and experimental species concentrations over time using a version of the BFGS algorithm. At each optimization step, the initial value problem was solved numerically for a given set of rate constants, and the resulting time-dependent concentrations were sampled at the corresponding experimental time points. All kinetic fitting and simulations were performed using the SciPy Python package, as well as code developed in-house, which is available from github.com/refeyn/publications/tree/master/ivonescimab-kinetics.

## Results

### Mass photometry profiles of ivonescimab, VEGF and PD-1

We first characterized the interaction of ivonescimab with its targets separately, before investigating their dynamics when mixed. This approach is good practice for binding studies with MP, as it on the one hand is a quality control step to ensure the components have not degraded or aggregated, and on the other hand it facilitates peak assignment and interpretation of the results. We first measured ivonescimab in isolation. Based on its primary sequence, we expected the molecular mass of ivonescimab to be 201 kDa. MP measurements of ivonescimab showed monomers were the predominant species in solution with a measured mass of 208 kDa (Fig. 1A). Dimers, which had an expected molecular mass of 416 kDa and a mass of 419 kDa as measured by MP, represented 11% of the total detected particles (Fig. 1A). Trimers, which had an expected molecular mass of 624 kDa and a mass of 626 kDa as measured by MP, represented ≈2% of the particles (Fig. 1A). This analysis indicated that the MP mass measurements of ivonescimab monomers and oligomers were in excellent agreement with the expected mass values.

**Fig. 1.**
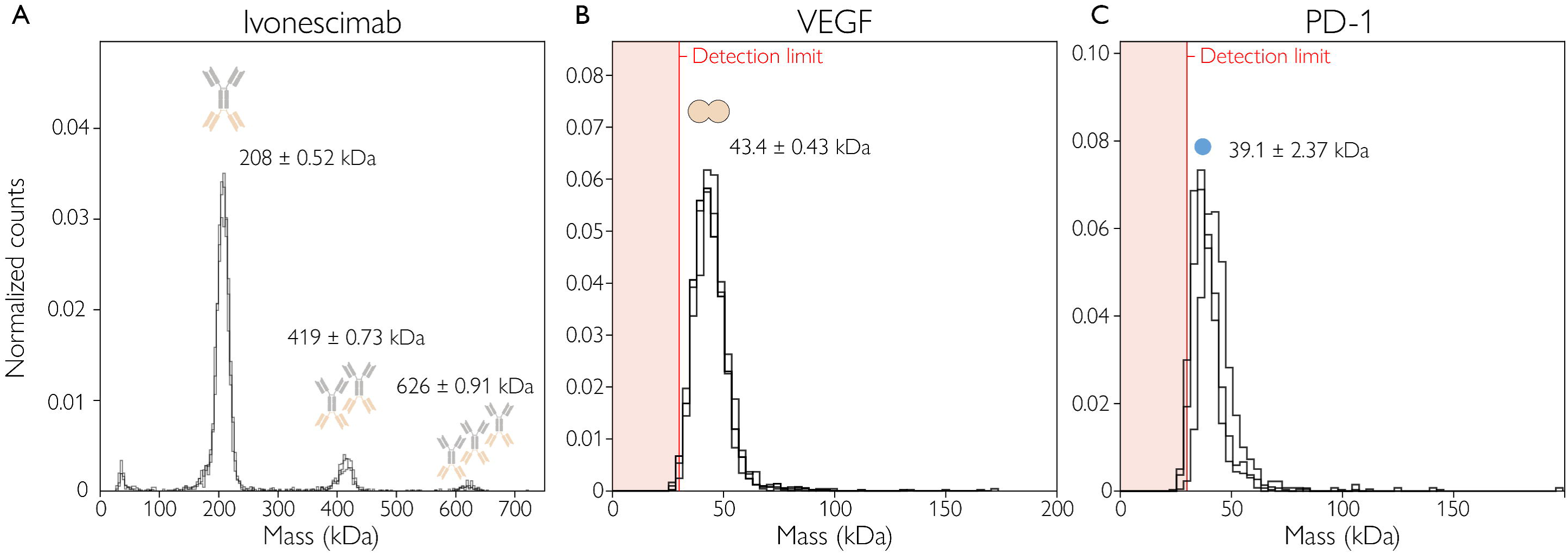
Mass photometry profiles of (A) ivonescimab and (B) VEGF and (C) PD-1. Each plot shows the counts (normalized) of particles detected within each mass bin vs. the molecular mass. Sample concentrations were 2.3–3.9 nM for ivonescimab, 1.8 nM for VEGF and 2 nM for PD-1. Measurements were performed in triplicate. For all measurements, the labels indicate the mean mass value ± standard deviation (SD) of the triplicate measurements and the schematics above each peak represent putative complex structures corresponding to those peaks. Red vertical line indicates the instrument’s lower limit of detection (30 kDa); there may be species present in the shaded range (0–30 kDa) that were not detected.

We compared this aggregation behavior to that of two other anti-PD-1 antibodies: penpulimab and pembrolizumab (commercialized as Keytruda). Both antibodies had high proportions of monomers. Penpulimab existed only as a 149 kDa monomer; in the case of pembrozilumab, dimers accounted for only 4% of the particles detected and trimers 1% (Fig. S1). This comparison confirmed that just like ivonescimab, other anti-PD-1 antibodies, also exist primarily as monomers.

Next, we investigated each of the ivonescimab targets, VEGF and PD-1, in isolation. VEGF is a homodimer, with an expected mass between 40 and 45 kDa according to the supplier. Our MP measurements were again in close agreement with the expected mass, showing a single peak at 43 kDa (Fig. 1B).

For PD-1, on the other hand, the picture is more complex, likely due to glycosylation and other modifications. The anti-PD-1 binding site of ivonescimab was constructed using Akeso Biopharma’s PD-1 antibody penpulimab (AK105),^7^ so the epitope for ivonescimab binding is the highly glycosylated N58 residue in PD-1, as shown by structural studies.^15^ Human PD-1 is known to be a highly glycosylated protein, with four major glycosylation sites (N49, N58, N74, and N116). Further modifications like fucosylation add complexity to the glycosylation at N58 and can significantly impact binding affinities for antibodies targeting the N58 epitope.^16^ According to SEC-MALS data from the supplier, the mass of PD-1-His has the range 25-40 kDa, indicating clear heterogeneity in the sample. This result was confirmed with SDS-PAGE analysis, also done by the supplier, which reported a mass range of 31-44 kDa. Further complicating the picture, glycomic profiling of PD-1 protein expressed in HEK cells has shown that not all N58 sites are occupied by glycans, indicating incomplete modification and high site-specific glycan microheterogeneity, meaning N58 on PD-1 carries multiple distinct glycoforms.^17^

When we used MP to measure PD-1, the mean mass value was 39.1 kDa (Fig. 1C) – consistent with the ranges reported from SEC-MALS and SDS-PAGE measurements. However, because the lower limit of detection for MP is 30 kDa, PD-1 molecules with mass at the lower end of the range may have been undetected (Fig. 1C, red line), which would cause the mean measurement for the population to be skewed to higher values and the amount of PD-1 present to be underestimated.

### Ivonescimab and VEGF interaction dynamics and stoichiometry

Next, we investigated the interaction of ivonescimab with VEGF. Specifically, we focused on determining which binding stoichiometries and which higher-order complexes of VEGF and ivonescimab molecules would be the most abundant, which has not been reported thus far. First, to observe the real-time dynamics of ivonescimab-VEGF binding, ivonescimab was incubated with VEGF in a 1:2 ratio, and the mixture was measured by MP at different time points. For each measurement, the mixture was incubated for the time indicated (Fig. 2A) then quickly diluted (1-3 µL in 20 µL of total PBS volume of droplet) to nanomolar concentration (as required for a standard MP measurement) before undergoing the measurement, which lasts 60 sec.

**Fig. 2.**
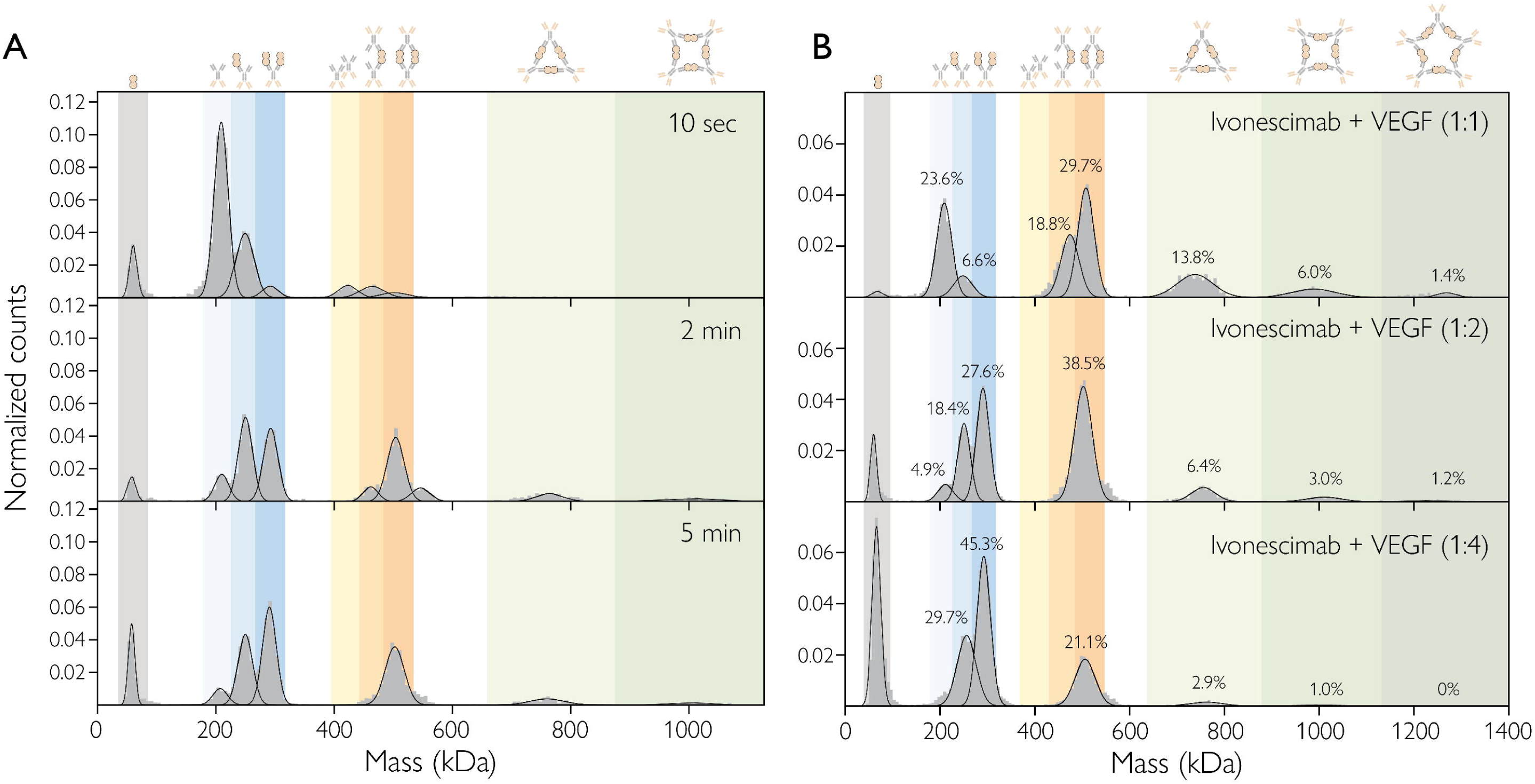
Mass photometry analysis of ivonescimab:VEGF interactions, over time and at equilibrium. (A) Ivonescimab:VEGF complex formation at 10 sec (top), 2 min (middle) and 5 min (bottom) of incubation of the sample mixture. Samples were incubated at a 1:2 antibody:antigen molar ratio (antibody at 50 nM) and imaged with the large field of view setting. The schematics above each peak represent putative complex structures. (B) Ivonescimab:VEGF complex formation for mixtures with 1:1, 1:2 and 1:4 antibody:antigen concentration ratios, at equilibrium, after 40 min incubation. Immediately prior to measurement, samples were diluted 10x or 20x in a PBS droplet, resulting in a concentration of 2.5 or 5 nM (for the antibody). (A-B) Each plot shows the counts (normalized) of particles detected within each mass bin vs. the molecular mass. The schematics above each peak represent putative complex structures. The shading represents the mass ranges corresponding to the different species: single antigens (grey); single antibody (blue shades) or two antibodies (orange shades) with 0 (very light), 1 (light) or 2 (mid) antigens; three antibodies (light green) or four antibodies (mid green).

After 10 sec of incubation, in addition to peaks corresponding to free VEGF and ivonescimab, peaks corresponding to ivonescimab-VEGF complexes were visible. Based on their mass, these complexes contained: one ivonescimab molecule in complex with either one or two VEGF molecules (249 kDa, 292 kDa), and complexes containing two ivonescimab molecules alone (423 kDa) or in complex with either one or two VEGF molecules (468 kDa, 509 kDa) (Fig. 2A, top). At this timepoint, the predominant species was free ivonescimab and the most abundant complex was 1:1 ivonescimab:VEGF (Fig. 2A, top).

After 2 minutes of incubation, the composition of the mixture changed dramatically. The three largest peaks all corresponded to complexes: one ivonescimab bound to either one or two VEGF molecules as well as 2:2 ivonescimab:VEGF complexes. Small quantities of 3:3, 4:4 and 5:5 ivonescimab:VEGF complexes also appeared, with masses of 761 kDa, 1015 kDa and 1269 kDa, respectively. Meanwhile, the abundance of free ivonescimab decreased considerably; it accounted for 58% of the particles after 10 sec of incubation but just 9% after 2 min. This shift continued over the next few minutes, with the measurement after 5 min of incubation showing a further reduction in the abundance of free ivonescimab to 6%, along with further slight increases in the abundance of the 1:2 and 2:2 complexes. From 5 min of incubation onward, no further changes were observed, leading us to conclude that the reaction reached equilibrium after 5 min of incubation (not shown).

Once we had established the time it takes to reach equilibrium, we proceeded to investigate the effects of different ivonescimab:VEGF ratios on the reaction equilibrium – to better understand which stoichiometries are favored in each and gain insights into assembly pathways. Under the 1:2 antibody:antigen concentration ratio (Fig. 2A), the most abundant species at equilibrium were 1:2 and 2:2 ivonescimab:VEGF complexes. When ivonescimab was incubated with VEGF in a 1:4 ratio, however, free ivonescimab was depleted as 1:2 ivonescimab:VEGF complexes formed – limiting the assembly of higher-order complexes (Fig. 2B bottom). When the antibody:antigen ratio was 1:1, making free ivonescimab abundant, the equilibrium of the reaction shifted towards the 2:2 complexes (Fig. 2B top). These observations are consistent with an assembly pathway for 2:2 complexes that involves free ivonescimab binding to 1:2 complexes (as opposed to two 1:1 complexes coming together to form a 2:2 complex). One important implication is then that a critical minimal concentration of ivonescimab must be present in the TME in order to drive formation of the higher-order complexes that likely give rise to the higher apparent affinity of ivonescimab for VEGF^15^.

### Calculation of K_D_ for ivonescimab-VEGF complexes

The dimeric nature of VEGF and multivalency of ivonescimab result in an equilibrium of simple, crosslinked, and multimeric species with distinct affinities at each stage. An advantage of using MP is that it enables us to quantify the abundance of each species that can be resolved, from which we can determine the affinities of each interaction, provided that the affinities are in the nanomolar range. We used the equilibrium MP data with all species present (Fig. 2B), to build an assembly model that we used to calculate the dissociation constants (K_D_) of the different complexes, following a method described previously^14,18^.

The calculated K_D_ values are presented in Fig. 3 with illustrations of putative assembly structures. We found that the formation of the 2:2 ivonescimab:VEGF complex was highly favored with a K_D_ of 0.08 nM. All other complexes had binding affinities at least an order of magnitude higher (3.17 nM), suggesting that the additional steps that result in higher-order structures may be less favorable due to steric hindrance or entropic costs. The stepwise equilibrium model describes sequential assembly of the multivalent complexes. The K_D_ for the 1:1 interaction corresponds to the microscopic binding event, whereas the higher-order stepwise constants are apparent/macroscopic values that incorporate symmetry and statistical degeneracy effects arising from multivalent assembly.

**Fig. 3.**
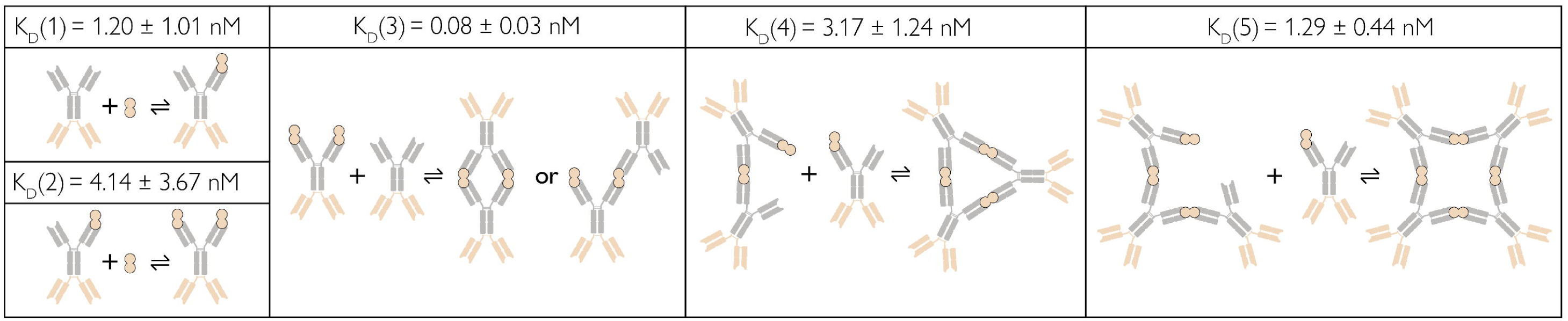
**Proposed ivonescimab**–**VEGF complex assembly models, with affinity quantification calculated from MP data.** Assembly models for the ivonescimab–VEGF complexes and calculated K_D_ values for reactions leading to the formation of ivonescimab–VEGF complexes with stoichiometries of (from left) 1:1 (upper) or 1:2 (lower), 2:2, 3:3 and 4:4. The K_D_ values were calculated from the MP data shown in Fig. 2, as described in the Methods.

We theorize that assembly starts with antibody and VEGF binding, then proceeds through secondary ligation events, and that it is dominated by strong crosslinking, where two antibodies are bridged by two VEGF dimers. As illustrated in Fig. 3, we predict that at each assembly stage, after a certain incubation time and depending on the molar ratios, free antibody and VEGF can participate in multiple pathways due to the multivalency of both components, resulting in multiple complex stoichiometries, of which the 2:2 complex is most abundant.

Dissociation constants, calculated from the abundances of the reactants and products at equilibrium (measured via MP single-molecule counts), were determined. They were similar for the ivonescimab-VEGF binding that results in the 1:1 complex, and then the additional VEGF binding that forms the 1:2 complex. K_D_ = 1.20 nM for the 1:1 complex, and K_D_= 4.14 nM for the 1:2 ivonescimab-VEGF complex (Fig. 3). These values are in excellent agreement with reported K_D_ values measured for the affinity of VEGF and ivonescimab using BLI (K_D_ =1.96 nM), when ivonescimab was immobilized on the surface^7^. However, a limitation of BLI is that bivalent interactions are not always easily detected, so this K_D_ value does not necessarily correspond to each specific, stepwise binding interaction (as do the K_D_ values calculated from the MP data).

The suggested mechanism of action described for ivonescimab is based on the observation that the formation of soluble ivonescimab-VEGF complexes, driven by VEGF, promotes binding between ivonescimab and human PD-1, mainly due to a lower dissociation rate^7^. The published data show the presence of higher-order ivonescimab-VEGF complexes, but no further details on dynamics, stoichiometry and equilibrium distribution. We show that, based on the MP data, ivonescimab-VEGF complexes with at least five different stoichiometries are present, all with different abundances. Even more striking are the apparent differences in dissociation constants. As discussed before, the VEGF binding steps that formed the 1:1 and 1:2 complexes showed similar K_D_s. The very low K_D_ value for the 2:2 stoichiometry (0.08 nM, 15x lower than the K_D_ for the 1:1 complex and 51x lower than the 1:2 complex) indicates a significantly tighter interaction. K_D_ values calculated for 3:3 and 4:4 complexes (3.17 nM and 1.29 nM) were higher than that for the 2:2 stoichiometry and similar to the values for the 1:1 and 1:2 complexes, indicating a weaker interaction. Accordingly, the 2:2 complex was most abundant in conditions with excess VEGF, while the abundances of 3:3, 4:4 and 5:5 complexes were significantly lower, with abundance decreasing as the size of the structure increased.

Although our data do not provide direct insights into assembly structure, we have provided putative structures to aid understanding of the components present and interactions (Fig. 3), which are based on findings in a recent study of complexes that form with the IL-17A cytokine (a homodimer) and anti-IL-17A antibodies^12^. There, exploring target-related immune complexes, which can trigger the formation of antibody-drug antibodies, the authors used SEC, AUC, and MP to show that a 2:2 stoichiometry was dominant for three of the four antibodies they studied. They also used negative-stain transmission electron microscopy (nsTEM) to confirm formation of closed-chain structures for all four antibodies – each involving at least 2 IgG molecules and an equal number of IL-17A dimers^12^. The dominance of the 2:2 complex that we observed is also consistent with observations of a 2 VEGF:Fab structure in a previous study^19^.

### Dynamics, stoichiometry and kinetic analysis of the ivonescimab – PD-1 interaction

Next, we examined the interactions between ivonescimab and PD-1, using a similar procedure to the one we used to study the ivonescimab:VEGF interactions (see previous section). As before, we incubated ivonescimab and PD-1 antigen in a 1:2 concentration ratio (with ∼10 nM ivonescimab), and used MP to measure the sample mixture at different time points (Fig. 4A). Equilibrium was reached within 10 min. Initially evident (after 10 sec incubation) were peaks corresponding to free PD-1 antigen, free antibody (212 kDa), a small population of complexes with 1:1 stoichiometry (242 kDa) and a small population of antibody dimers (438 kDa).

**Fig. 4.**
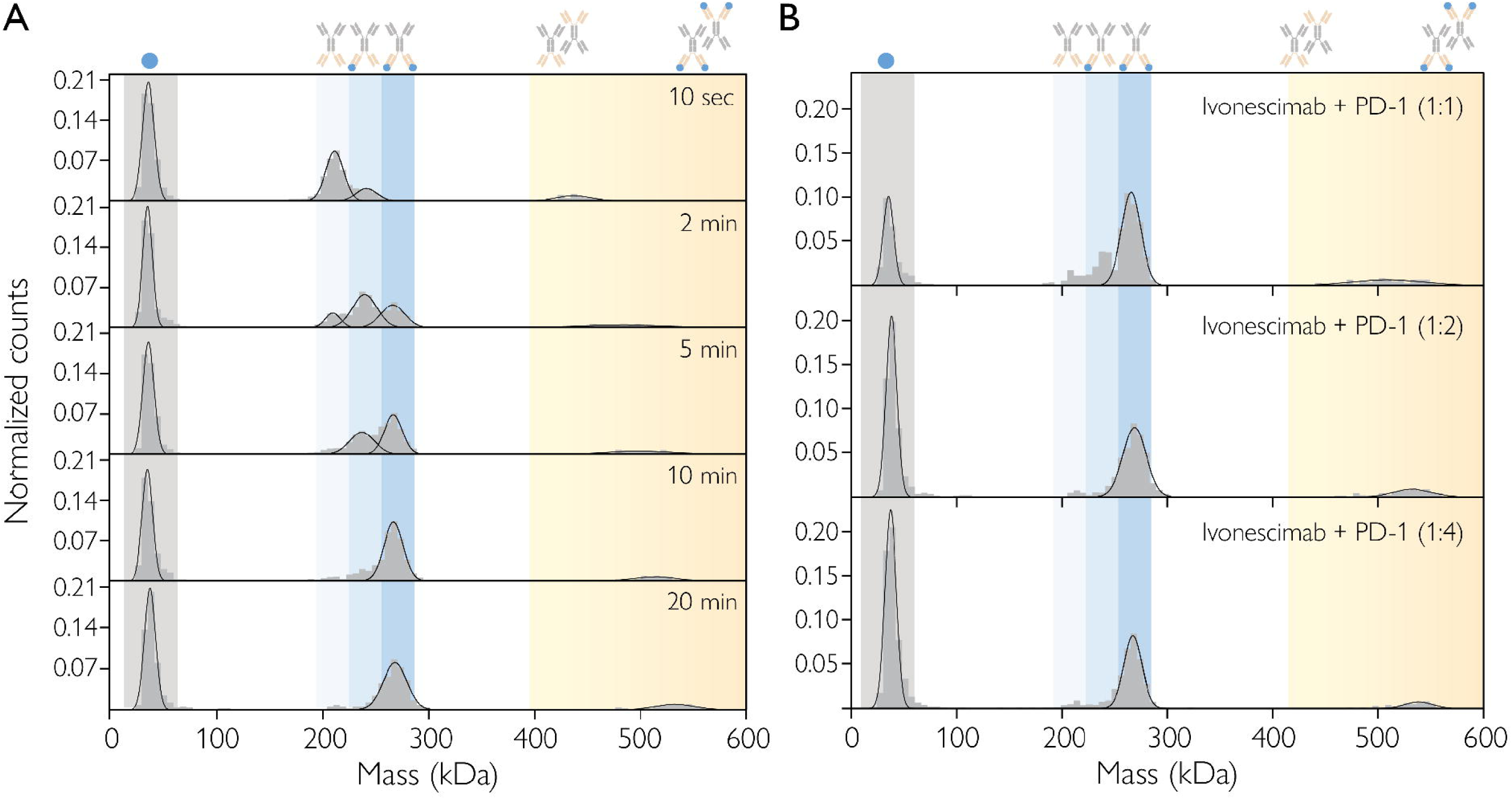
MP analysis of ivonescimab and PD-1 interactions, over time and at equilibrium. (A) Ivonescimab–PD-1 complex formation at different time points of incubation of the sample mixture. Samples were incubated at a 1:2 antibody:antigen molar ratio (antibody at 10 nM). Imaged in regular field of view. The schematics above each peak represent putative structures for each complex. (B) Ivonescimab–PD-1 complex formation at equilibrium, after 40 min incubation, for antibody:antigen molar ratios of 1:1, 1:2 and 1:4 (antibody at 10 nM). Prior to measurement, samples were diluted 4x in a PBS droplet, resulting in 2.5 nM antibody conc. Each plot shows the counts (normalized) of particles detected within each mass bin vs. the molecular mass. The schematics above each peak represent putative complex structures. The shading represents the mass ranges corresponding to the different species: single antigens (grey); single antibody (blue shades) with 0 (very light), 1 (light) or 2 (mid) antigens; or two antibodies with 0-2 antigens (orange).

When measuring ivonescimab-PD-1 complexes, we observed that the mass shift between ivonescimab alone and bound to PD-1 was around 30 kDa – indicating that PD-1 has a mass of 30 kDa. This is within the range of values measured by SEC-MALS and SDS-PAGE. However, it was surprising because the mass of PD-1 that we measured with MP for PD-1 in isolation was 39.1 kDa. One possible explanation for this discrepancy is that, as mentioned above, PD-1 appears highly heterogeneous, likely due to modifications such as glycosylation. It may be that only a subset of all the PD-1 present is in the correct form to bind to ivonescimab, and it has a mass of around 30 kDa. In support of this hypothesis, the antibody penpulimab (which was used in the design of ivonescimab) is known to bind preferentially to a specific glycosylated form of PD-1^15^. Another possible explanation is that the actual mean mass of the PD-1 present is lower than the measured value of 39.1 kDa, but MP has failed to detect lower-mass PD-1 species (due to its lower limit of detection being 30 kDa), so the mean is skewed to a higher mass value. In either case, it means that – unlike for the other species – the effective abundance of PD-1 present in the mixtures measured may not reliably correspond to the measured counts. That is, the counts may be an overestimate (if some observed molecules are not ‘available’ to interact with ivonescimab due to their glycosylation status) or they may be an underestimate (if only some of the molecules were detected). We factor in this uncertainty in our further analyses involving PD-1.

Over time, the amount of free antibody present decreased; after 10 or 20 min of incubation, almost all ivonescimab was in complex with PD-1. Meanwhile, after 2-4 min of incubation, populations of ivonescimab–PD-1 complexes with stoichiometries of both 1:1 (241 kDa) and 1:2 (269 kDa) were apparent (Fig. 4A). The 1:1 complex was transient, however, and had mostly disappeared by 10-20 min of incubation, indicating that the 1:2 complexes are the more stable assembly. As the system approached equilibrium, another small population (539 kDa) with mass consistent with two ivonescimab (418 kDa) and four PD-1 (∼120 kDa) molecules was also detected. Based on its mass, we conclude that this population is ivonescimab dimers with one PD-1 bound to each of the four available binding sites. It accounted for ∼10% of the total counts.

We also measured samples with antibody:antigen ratios of 1:1 and 1:4 (with 10 nM ivonescimab). We found that the incubation time required to reach equilibrium varied significantly from 40 min for a 1:1 sample to only 5 min at a 1:4 ratio (Suppl. Fig. 2). This variability is likely due to the availability of PD-1 combined with the high-affinity interaction between ivonescimab and PD-1. When the antigen is in excess, the favored complexes (with a 1:2 stoichiometry) can readily form; when there is less antigen, a 1:1 complex is also detectable.

Another broad peak could also be seen around 510 kDa in the 1:1 sample (Fig. 4B), and its mean mass corresponds to two ivonescimab molecules and three PD-1 antigens. Because the peak is quite broad, there are likely to be multiple ivonescimab dimer species present, in complex with 1 to 4 molecules of PD-1. In the other samples, where more PD-1 is present (1:2 and 1:4 ratios), this complex disappeared in favor of the 2:4 complex described above (∼539 kDa).

When the ivonescimab–PD-1 interaction was at equilibrium, at molar ratios of 1:1, 1:2 and 1:4 (Fig. 4B), ivonescimab was fully saturated by PD-1, existing only in the ivonescimab:(PD-1)_2_ complex. Since no free ivonescimab was present, we could not calculate the K_D_ from the equilibrium data, as we did for the ivonescimab-VEGF interaction. Consequently, we instead used real-time data for the formation of the ivonescimab:PD-1 and ivonescimab(PD-1)_2_ complexes (Fig. 5A), where we mixed ivonescimab and PD-1 at a 1:4 molar ratio with a 5 nM antibody concentration, without further droplet dilution, to fit a mathematical model of the reaction using data from three repeat time series of a 1:4 mix of ivonescimab and PD-1, in order to extract kinetic parameters. We chose to model PD-1 concentrations using mass balance due to our uncertainty about its abundance, as discussed above. We modelled the reaction, using ordinary differential equations, as a two-step process where first one molecule of free PD-1 binds to free ivonescimab and then a second free PD-1 binds to an ivonescimab–PD-1 complex (Fig. 5A). Rates of complex formation appeared similar for both ivonescimab monomers and dimers binding to PD-1 (Fig. 4). Specifically, the dimer peak (10% of total ivonescimab) disappeared with similar kinetics to the monomer peak, while the dimer:4PD-1 appeared in parallel. From this we concluded that Ivonescimab dimers bind to PD1 with similar affinity to monomer under the given kinetic assay conditions.

**Fig. 5.**
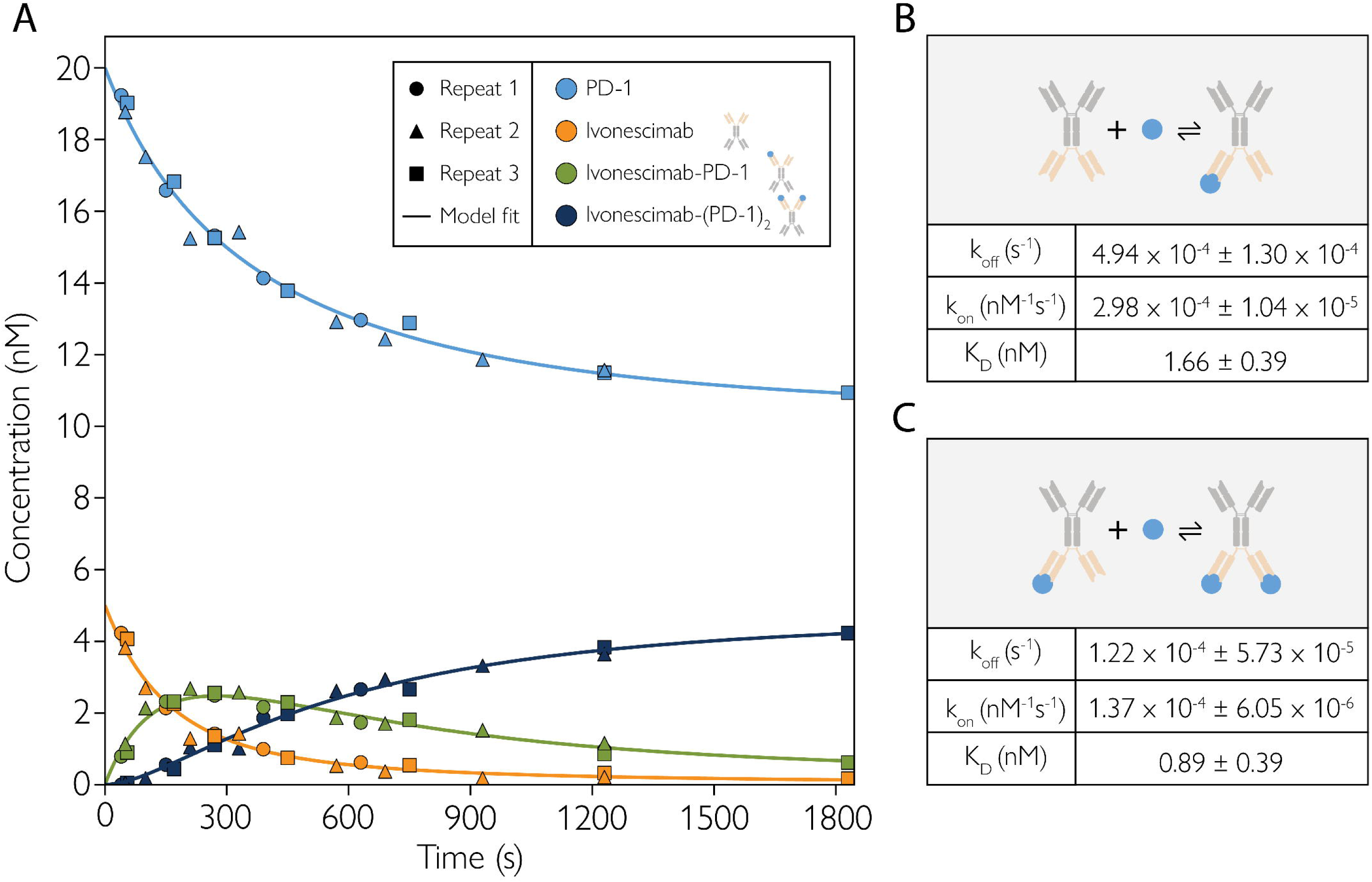
Quantitative modelling of ivonescimab binding to PD-1 reveals formation dynamics and affinities for complexes with distinct stoichiometries. (A) Time course of ivonescimab and PD-1 complex formation at 5 nM ivonescimab and initial PD-1 concentration of 20 nM, showing model simulation (lines) and MP data (shapes) for an experiment performed in triplicate. The concentrations of free PD-1 (blue), free ivonescimab (orange), ivonescimab-PD-1 (green), and ivonescimab-(PD-1)_2_ (black) are plotted over time. Fitting was done as described in the Materials and Methods. (B,C) Kinetic and affinity parameters for the sequential steps of PD-1 binding to ivonescimab – the binding of the first (B) and second (C) PD-1 molecules. The SD values are derived from the Jacobian (covariance) matrix at the least squares optimum and the K_D_ values were calculated as k_off_ /k_on_.

A model simulation is shown alongside the repeat measurements (Fig. 5A), demonstrating the rapid initial depletion of free ivonescimab species and subsequent accumulation of fully saturated ivonescimab-PD-1_2_ complex at equilibrium. The two sequential binding events display similar affinities (K_D_) in the low nanomolar range; 1.66 nM and 0.89 nM, with dissociation rates (*k_off_*) of 4.94E-4 s^-1^ and 1.22E-4 s^-1^ (Fig. 5B-C). These values are in agreement with prior measurements of this system done using BLI (where the measured values were K_D_= 0.72 nM and *k_off_* =2.10E-4 s^-^1)^7^, but, as for the ivonescimab-VEGF measurements, it is not clear whether a monovalent or bivalent model was used for fitting.

As described in the Methods, the empirical concentration of PD-1 (20 nM) was assigned as the total PD-1 concentration in the model fitting, due to uncertainty about the reliability of the MP measurements of free PD-1. It is likely that only some PD-1 is accessible to ivonescimab for binding (due to not all being in the same state of glycosylation), making 20 nM an overestimate. If the true (or effective) PD-1 concentration was indeed slightly lower, this would result in slightly lower K_D_ values (Table S1).

### Ivonescimab oligomerization is driven by VEGF and not affected by PD-1

Having investigated the interaction of ivonescimab with its two targets separately, we moved on to analyze the formation of ivonescimab–VEGF–PD-1 complexes. First, ivonescimab was incubated with VEGF in a 1:2 ratio. As before, equilibrium was established after 40 min and the ivonescimab–VEGF complexes formed were consistent with those shown in Fig. 2 (Fig. 6A, top). Next, PD-1 was added for a final 1:2:2 ratio and complex formation was monitored over time. Equilibrium was reached after 40 min (data not shown). The number and overall distribution of peaks did not change after the addition of PD-1, but they did increase in mass (Fig. 6A, bottom, detailed in Fig. 6B). For each peak, this mass increase was consistent with the addition of two PD-1 molecules for each ivonescimab molecule in the complex. The mass distribution of the sample did not change when the order of addition of components was changed (Fig. S3). More specifically, when ivonescimab was preincubated with PD-1 (Fig. S2 right lower panel), addition of VEGF drove the formation of the same higher-order structures with the same mass distribution, in a similar time period as when ivonescimab was incubated with VEGF only, or with VEGF and PD-1 together. The results show that VEGF drives higher-order complex formation but does not alter the stoichiometry of PD-1 binding to ivonescimab. They also show that PD-1 does not promote or disrupt ivonescimab oligomerization, which is driven by VEGF binding. Whether there is cooperativity, i.e. PD-1 binds with higher affinity to ivonescimab if VEGF is bound to it, as suggested in an earlier study,^7^ was not assessed here.

**Fig. 6.**
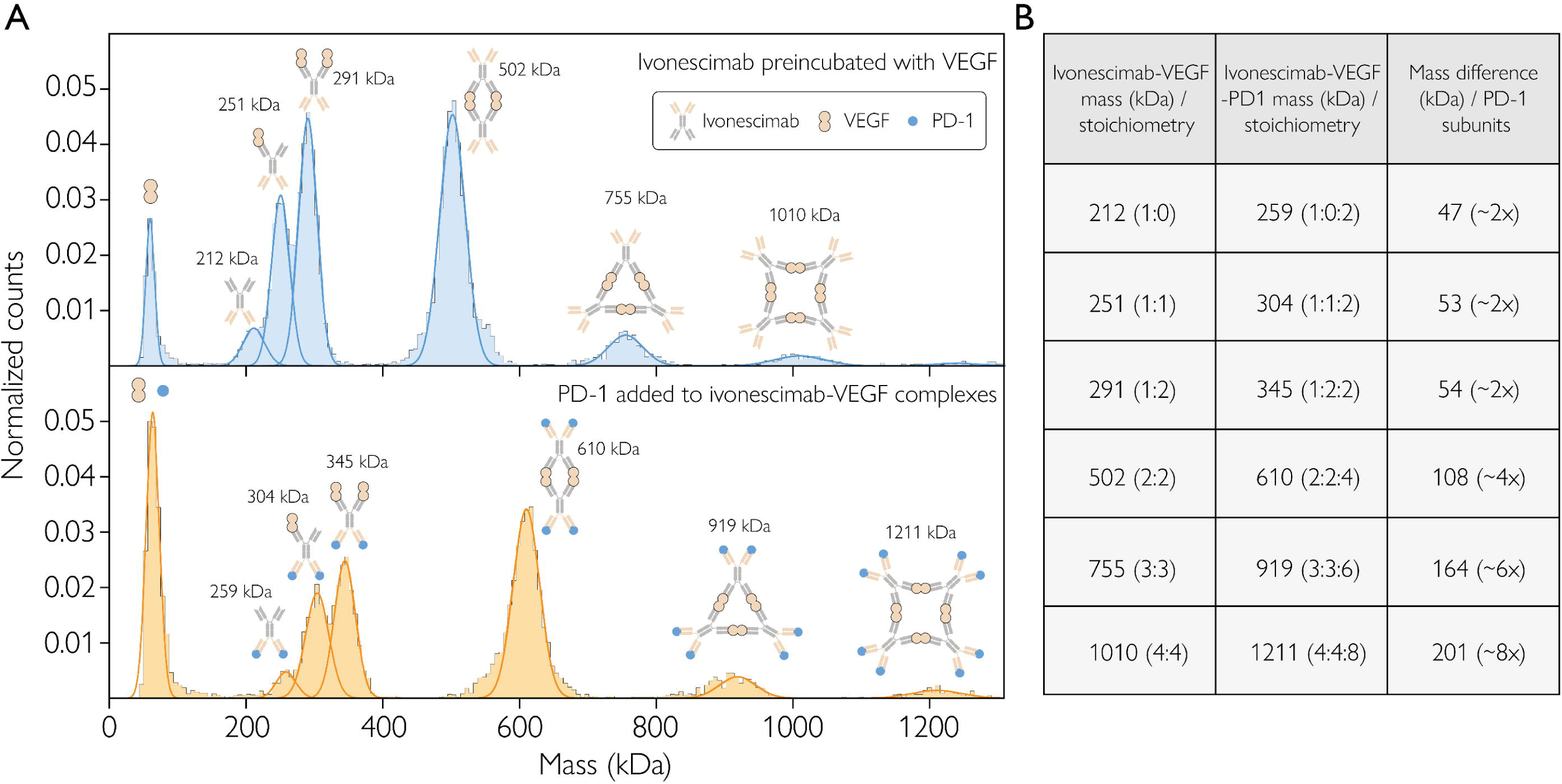
Mass photometry analysis of Ivonescimab, VEGF, and PD-1 complex formation. (A) Top: Ivonescimab–VEGF complex formation at equilibrium, with a 1:2 concentration ratio. Bottom: Ivonescimab–VEGF–PD-1 complex formation at equilibrium, with a 1:2:2 mixing ratio. Each plot shows the counts (normalized) of particles detected within each mass bin vs. the molecular mass. The schematics above each peak represent putative complex structures. (B) Overview of all the complexes detected in A (top) compared to the ones detected in A (bottom), quantifying the mass shift caused by the addition of PD-1.

## Discussion

In this paper, we used MP to characterize ivonescimab, a next-generation bispecific antibody, and its binding to both of its targets, PD-1 and VEGF, *in vitro*. Although *in vitro* characterization of ivonescimab has been attempted before, it had not been possible to obtain a detailed picture of the binding dynamics and the stoichiometry of the complexes formed. Using the single-molecule mass measurement technique MP, we were able to characterize the formation and stoichiometries of these complexes, at near-physiological concentrations. Although concentrations within the TME are not fully known, there is evidence, for example, that the concentration of VEGF is on the order of pg/mL in human plasma^20^, but is elevated to nM concentrations in cancer patients^21^. As PD-1 is on the cell surface, physiological concentrations are more difficult to estimate. Most studies report relative PD-1 expression, comparing high expression levels on activated T cells in the TME to low levels in non-activated/peripheral T cells^22^. By measuring samples containing mixtures of ivonescimab and either VEGF or PD-1 at different time points, before and after equilibrium had been reached, we were able to track complex formation between ivonescimab and its targets and calculate the affinities of key interactions. Our results provide new insights into the mechanism, stoichiometry and dynamics of complex formation for ivonescimab and its targets.

It was previously thought that VEGF binds to ivonescimab, driving the formation of large oligomers^7^. This hypothesis builds on the observation that the kinetic off-rate for the interaction of PD-1 with ivonescimab decreased 18-fold when VEGF was present, though it was unclear whether this could be an avidity or cooperativity effect.

We have confirmed that VEGF drives oligomerization of ivonescimab, which appears primarily as a monomer on its own, similar to other anti-PD-1 antibodies. However, we have also shown that the most stable structure formed by these two molecules is a 2:2 complex, not the expected higher-order oligomers. This finding is significant as it was not apparent from prior studies with other methods, and the stoichiometry directly impacts the interpretation of downstream bioanalytical and pharmacokinetic assays. Another study examining antigen–antibody structures of IL-17A (which has a cystine-knot structure like VEGF) with four different anti-IL-17A antibodies reported that three of the antibodies formed 2:2 antigen–antibody complexes^12^. This indicated that the 2:2 complex is a stable, closed-loop structure, such that dissociation requires disruption of multiple Fab–antigen interactions, reflecting an avidity effect that stabilizes the overall complex. A similar mechanism could play a role in VEGF-ivonescimab interactions.

Considering these interactions in more depth, we observe that, for VEGF-ivonescimab binding, K_D_(1)=1.20 nM for the first binding site and K_D_(2)=4.14 nM for the second – and the slightly higher value of K_D_(2) makes sense as, for two equivalent binding sites, binding to the second site would have an on-rate of half the first due to the first site being occupied. As mentioned above, these values are in excellent agreement with BLI results (K_D_ =1.96 nM) when ivonescimab was immobilized on the surface^7^. However a different K_D_ value (K_D_ =0.33 nM) was obtained when VEGF was immobilized – underlining the value of a solution-based approach in avoiding avidity effects. The formation of the 2:2 complex had the highest affinity by far (K_D_(3) = 0.08 nM), which may suggest that two ivonescimab-VEGF interactions are occurring at once, resulting in a strong enthalpic drive to form this complex. The K_D_(3) value reflects a fully bridged, native, high-avidity complex in solution, formed without surface constraints or ligand-density-driven avidity and rebinding, which are common issues with BLI and SPR. Additionally, even if one of the interactions is broken, the molecule remains tethered with the free site in close proximity, facilitating reattachment. Looking at K_D_(4) (3.17 nM), the higher value reflects the need for sufficient free ivonescimab to be present to continue forming the chain (and this needs to outcompete formation of the stable 2:2 structure). Overall, higher-order structures do not appear to be favored, as evidenced by the low prevalence of 3:3, 4:4, and 5:5 complexes and their correspondingly higher K_D_ values.

Our analysis of the interaction between ivonescimab and PD-1 was complicated by the heterogeneity of PD-1 – likely to due to glycosylation or other modifications – and by the mass range of the PD-1 species (25 – 40 kDa) overlapping the lower limit of detection of the mass photometer (30 kDa). This meant that our mass photometry measurements of the abundance of free PD-1 may not have been accurate, which we overcame by assigning the empirical concentration used as its initial concentration, then using the principle of mass balance to calculate its concentration over time, as it entered into complexes with ivonescimab that could be reliable detected and quantified. With this approach, and using our kinetic model constrained by real-time MP measurements, we found that the two sequential binding events, for the binding of the first and second PD-1 antigens to ivonescimab had similar affinities (K_D_) of 1.66nM and 0.89 nM, in agreement with prior BLI measurements^7^. We also tracked real-time complex formation between ivonescimab and both its targets at the same time. Interestingly, the complexes that formed when all three molecules were present followed the same distribution as seen in mixtures of ivonescimab and VEGF alone, with PD-1 consistently being added in a 2:1 ratio, resulting in the 2:2:4 complex (of ivonescimab:VEGF:PD-1) being most abundant. This suggests that PD-1 does not promote or inhibit further oligomerization, or otherwise have an effect elsewhere in the complex when it binds to ivonescimab.

Another interesting finding of this study is that higher-order structure formation of ivonescimab with VEGF depends on the molar ratio between the two molecules. If there is more VEGF than antibody, the binding sites get saturated and higher-order structures are less likely to form. Hence, MP was used to determine that there is a critical amount of ivonescimab required in the TME for oligomerization to happen and the therapeutically beneficial higher-order structures to form. This means that MP is an important technique in any dosage studies where complex formation is a critical factor, as it informs about the concentration ratio required to achieve the optimal local concentration of the pharmaceutically active complex.

These results advance our understanding of ivonescimab-VEGF interactions and highlight the value of MP for this type of investigation. Identifying the stoichiometry for this clinically relevant combination of antibody and antigens is important, as is the demonstration of a reliable approach to obtain this information for antibody-antigen systems. One advantage of MP is that it allows complex formation to occur in solution. Both ivonescimab and VEGF are soluble in the TME, so the conditions here were closer to physiological conditions for this type of complex formation. In contrast, surface-based methods like SPR and BLI, due to the immobilization of components, can result in either the blocking of binding sites or false avidity (if the immobilized VEGF is not distributed sparsely enough); they can also obscure solution-phase oligomerization, rebinding, and assembly pathways. Revealing complex formation and association/dissociation kinetics in near-physiological conditions (label-free and in solution) enables more precise protein engineering, to achieve better binding and stability. In addition, while techniques like SPR and BLI are ideal for monovalent binding, they require more optimization and fitting to describe bivalent interactions, which are otherwise not always easily detected from SPR and BLI data.

Understanding the binding dynamics and stoichiometry of ivonescimab’s interactions with its targets, VEGF and PD-1, is an important step towards realizing this antibody’s therapeutic potential. The MP data presented here fills gaps in our prior understanding of these interactions, while demonstrating that MP complements techniques widely used during antibody discovery and development, like SPR and BLI. MP can determine which different stoichiometries are present and the relative abundance of each, supporting a more complete understanding of the action of multi-specific antibodies, and informing their design. More generally, MP-based approaches are a promising addition for research and development of new antibody modalities. MP clearly has the potential to support real-time monitoring of antibody quality to reduce bottlenecks and improve product consistency, from R&D through manufacturing scale-up.

## Acknowledgements

We thank Justin Benesch for helpful discussions, critical review of the manuscript, and valuable conceptual input. We also thank Nicolas Palanca for assistance with manuscript preparation.

## Data sharing

Kinetic model code developed in-house (and associated data) is available from github.com/refeyn/publications/tree/master/ivonescimab-kinetics/. Other data available upon request.

## Ethics and Consent

This study did not involve human or animal subjects or human data. All materials were commercially obtained and used in vitro. As such, ethical approval and informed consent were not required.

## References

(1) Matairi, A. A.; Hammadeh, B. M.; Aldalati, A. Y.; Qtaishat, F. A.; Nashwan, A. J.; Alzibdeh, A. Efficacy and Safety of Ivonescimab in the Treatment of Advanced Non-Small Cell Lung Cancer (NSCLC): A Systematic Review. Cureus 2025, 17 (1). 10.7759/cureus.77381.

(2) Khan, M.; Lin, J.; Liao, G.; Tian, Y.; Liang, Y.; Li, R.; Liu, M.; Yuan, Y. Comparative Analysis of Immune Checkpoint Inhibitors and Chemotherapy in the Treatment of Advanced Non-Small Cell Lung Cancer: A Meta-Analysis of Randomized Controlled Trials. Medicine (Baltimore) 2018, 97 (33), e11936. 10.1097/MD.0000000000011936.

(3) Sharma, P.; Hu-Lieskovan, S.; Wargo, J. A.; Ribas, A. Primary, Adaptive, and Acquired Resistance to Cancer Immunotherapy. Cell 2017, 168 (4), 707–723. 10.1016/j.cell.2017.01.017.

(4) Cui, J.-W.; Li, Y.; Yang, Y.; Yang, H.-K.; Dong, J.-M.; Xiao, Z.-H.; He, X.; Guo, J.- H.; Wang, R.-Q.; Dai, B.; Zhou, Z.-L. Tumor Immunotherapy Resistance: Revealing the Mechanism of PD-1 / PD-L1-Mediated Tumor Immune Escape. Biomed. Pharmacother. 2024, 171, 116203. 10.1016/j.biopha.2024.116203.

(5) Mahaki, H.; Nobari, S.; Tanzadehpanah, H.; Babaeizad, A.; Kazemzadeh, G.; Mehrabzadeh, M.; Valipour, A.; Yazdinezhad, N.; Manoochehri, H.; Yang, P.; Sheykhhasan, M. Targeting VEGF Signaling for Tumor Microenvironment Remodeling and Metastasis Inhibition: Therapeutic Strategies and Insights. Biomed. Pharmacother. 2025, 186, 118023. 10.1016/j.biopha.2025.118023.

(6) Saeed, A.; Park, R.; Sun, W. The Integration of Immune Checkpoint Inhibitors with VEGF Targeted Agents in Advanced Gastric and Gastroesophageal Adenocarcinoma: A Review on the Rationale and Results of Early Phase Trials. J. Hematol. Oncol.J Hematol Oncol 2021, 14 (1), 13. 10.1186/s13045-021-01034-0.

(7) Zhong, T.; Zhang, L.; Huang, Z.; Pang, X.; Jin, C.; Liu, W.; Du, J.; Yin, W.; Chen, N.; Min, J.; Xia, M.; Li, B. Design of a Fragment Crystallizable-Engineered Tetravalent Bispecific Antibody Targeting Programmed Cell Death-1 and Vascular Endothelial Growth Factor with Cooperative Biological Effects. iScience 2025, 28 (3). 10.1016/j.isci.2024.111722.

(8) Wang, F.; Wei, X.; Zheng, Y.; Wang, J.; Ying, J.; Chen, X.; Luo, S.; Luo, H.; Yu, X.; Chen, B.; Ma, L.; Xu, R. Safety, Pharmacokinetics, and Pharmacodynamics Evaluation of Ivonescimab, a Novel Bispecific Antibody Targeting PD-1 and VEGF, in Chinese Patients With Advanced Solid Tumors. Cancer Med. 2025, 14 (6), e70653. 10.1002/cam4.70653.

(9) Frentzas, S.; Austria Mislang, A. R.; Lemech, C.; Nagrial, A.; Underhill, C.; Wang, W.; Wang, Z. M.; Li, B.; Xia, Y.; Coward, J. I. G. Phase 1a Dose Escalation Study of Ivonescimab (AK112/SMT112), an Anti-PD-1/VEGF-A Bispecific Antibody, in Patients with Advanced Solid Tumors. J. Immunother. Cancer 2024, 12 (4), e008037. 10.1136/jitc-2023-008037.

(10) Xiong, A.; Wang, L.; Chen, J.; Wu, L.; Liu, B.; Yao, J.; Zhong, H.; Li, J.; Cheng, Y.; Sun, Y.; Ge, H.; Yao, J.; Shi, Q.; Zhou, M.; Chen, B.; Han, Z.; Wang, J.; Bu, Q.; Zhao, Y.; Chen, J.; Nie, L.; Li, G.; Li, X.; Yu, X.; Ji, Y.; Sun, D.; Ai, X.; Chu, Q.; Lin, Y.; Hao, J.; Huang, D.; Zhou, C.; Shan, J.; Yang, H.; Liu, X.; Wang, J.; Shang, Y.; Mei, X.; Yang, J.; Lu, D.; Hu, M.; Wang, Z. M.; Li, B.; Xia, M.; Zhou, C. Ivonescimab versus Pembrolizumab for PD-L1-Positive Non-Small Cell Lung Cancer (HARMONi-2): A Randomised, Double-Blind, Phase 3 Study in China. The Lancet 2025, 405 (10481), 839–849. 10.1016/S0140-6736(24)02722-3.

(11) Young, G.; Hundt, N.; Cole, D.; Fineberg, A.; Andrecka, J.; Tyler, A.; Olerinyova, A.; Ansari, A.; Marklund, E. G.; Collier, M. P.; Chandler, S. A.; Tkachenko, O.; Allen, J.; Crispin, M.; Billington, N.; Takagi, Y.; Sellers, J. R.; Eichmann, C.; Selenko, P.; Frey, L.; Riek, R.; Galpin, M. R.; Struwe, W. B.; Benesch, J. L. P.; Kukura, P. Quantitative Mass Imaging of Single Biological Macromolecules. Science 2018, 360 (6387), 423–427. 10.1126/science.aar5839.

(12) Ungan, D.; Be, C.; Baczyk, P.; Mittermeier, S.; Lehmann, S.; Wiesmann, C.; Huber, T.; Kolbinger, F.; Rondeau, J.-M. IL-17A Complexes with Therapeutic Antibodies Exhibit Distinct Size Distributions, Potentially Contributing to Clinically Observed Immunogenicity. mAbs 17 (1), 2575840. 10.1080/19420862.2025.2575840.

(13) Refeyn. TwoMP User Manual, V2.7, 2025.

(14) Soltermann, F.; Foley, E. D. B.; Pagnoni, V.; Galpin, M.; Benesch, J. L. P.; Kukura, P.; Struwe, W. B. Quantifying Protein–Protein Interactions by Molecular Counting with Mass Photometry. Angew. Chem. Int. Ed. 2020, 59 (27), 10774–10779. 10.1002/anie.202001578.

(15) Huang, Z.; Pang, X.; Zhong, T.; Qu, T.; Chen, N.; Ma, S.; He, X.; Xia, D.; Wang, M.; Xia, M.; Li, B. Penpulimab, an Fc-Engineered IgG1 Anti-PD-1 Antibody, With Improved Efficacy and Low Incidence of Immune-Related Adverse Events. Front. Immunol. 2022, 13, 924542. 10.3389/fimmu.2022.924542.

(16) Chu, C.-W.; Čaval, T.; Alisson-Silva, F.; Tankasala, A.; Guerrier, C.; Czerwieniec, G.; Läubli, H.; Schwarz, F. Variable PD-1 Glycosylation Modulates the Activity of Immune Checkpoint Inhibitors. Life Sci. Alliance 2024, 7 (3), e202302368. 10.26508/lsa.202302368.

(17) Izadi, S.; Abrantes, R.; Gumpelmair, S.; Kunnummel, V.; Duarte, H. O.; Steinberger, P.; Reis, C. A.; Castilho, A. An Engineered PD1-Fc Fusion Produced in N. Benthamiana Plants Efficiently Blocks PD1/PDL1 Interaction. Plant Cell Rep. 2025, 44 (4), 80. 10.1007/s00299-025-03475-0.

(18) Kofinova, Z.; Karunanithy, G.; Ferreira, A. S.; Struwe, W. B. Measuring Protein-Protein Interactions and Quantifying Their Dissociation Constants with Mass Photometry. Curr. Protoc. 2024, 4 (1), e962. 10.1002/cpz1.962.

(19) Walker, A.; Chung, C.-W.; Neu, M.; Burman, M.; Batuwangala, T.; Jones, G.; Tang, C.-M.; Steward, M.; Mullin, M.; Tournier, N.; Lewis, A.; Korczynska, J.; Chung, V.; Catchpole, I. Novel Interaction Mechanism of a Domain Antibody-Based Inhibitor of Human Vascular Endothelial Growth Factor with Greater Potency than Ranibizumab and Bevacizumab and Improved Capacity over Aflibercept *. J. Biol. Chem. 2016, 291 (11), 5500–5511. 10.1074/jbc.M115.691162.

(20) Kaya, A.; Çiledag, A.; Gulbay, B. E.; Poyraz, B. M.; Çelik, G.; Sen, E.; Savas, H.; Savas, I. The Prognostic Significance of Vascular Endothelial Growth Factor Levels in Sera of Non-Small Cell Lung Cancer Patients. Respir. Med. 2004, 98 (7), 632–636. 10.1016/j.rmed.2003.12.017.

(21) Raimondo, F.; Azzaro, M. P.; Palumbo, G. A.; Bagnato, S.; Stagno, F.; Giustolisi, G. M.; Cacciola, E.; Sortino, G.; Guglielmo, P.; Giustolisi, R. Elevated Vascular Endothelial Growth Factor (VEGF) Serum Levels in Idiopathic Myelofibrosis. Leukemia 2001, 15 (6), 976–980. 10.1038/sj.leu.2402124.

(22) Davidson, T. B.; Lee, A.; Hsu, M.; Sedighim, S.; Orpilla, J.; Treger, J.; Mastall, M.; Roesch, S.; Rapp, C.; Galvez, M.; Mochizuki, A.; Antonios, J.; Garcia, A.; Kotecha, N.; Bayless, N.; Nathanson, D.; Wang, A.; Everson, R.; Yong, W. H.; Cloughesy, T. F.; Liau, L. M.; Herold-Mende, C.; Prins, R. M. Expression of PD-1 by T Cells in Malignant Glioma Patients Reflects Exhaustion and Activation. Clin. Cancer Res. Off. J. Am. Assoc. Cancer Res. 2019, 25 (6), 1913–1922. 10.1158/1078-0432.CCR-18-1176.

