## Supplemental Material for "Mass photometry reveals stoichiometry and binding dynamics of bispecific tetravalent anti-VEGF-PD-1 antibody ivonescimab"

#### Supplementary Material

##### Figure S1

a) Pembrolizumab (Keytruda)

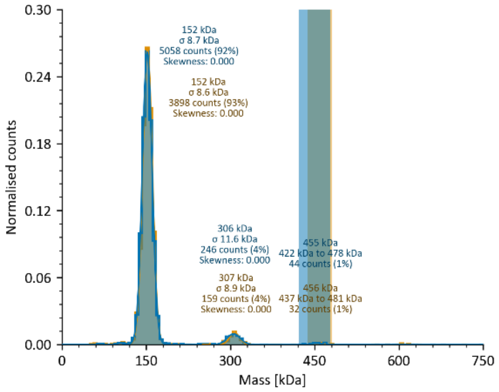


b) Penpulimab

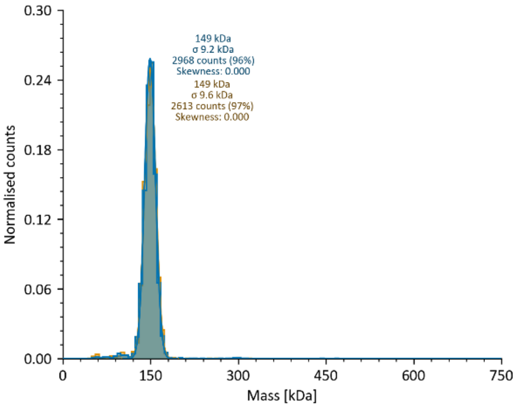


***Fig. S1 Mass photometry profiles of (A) pembrolizumab and (B) penpulimab.*** *The x-axis shows the measured molecular mass in kDa, while the y-axis shows the normalized counts for particles within each mass bin in the x-axis. Concentrations were 2.3-3.9 nM for ivonescimab and 1.8 nM for VEGF. The shaded regions around 450 kDa represent the user-defined mass range for trimers for each experiment. The labels report the mean and standard deviation for each fitted peak or region, as well as the corresponding counts and the peaks' skewness (asymmetry). Data shown are experimental duplicates.*

#### Figure S2

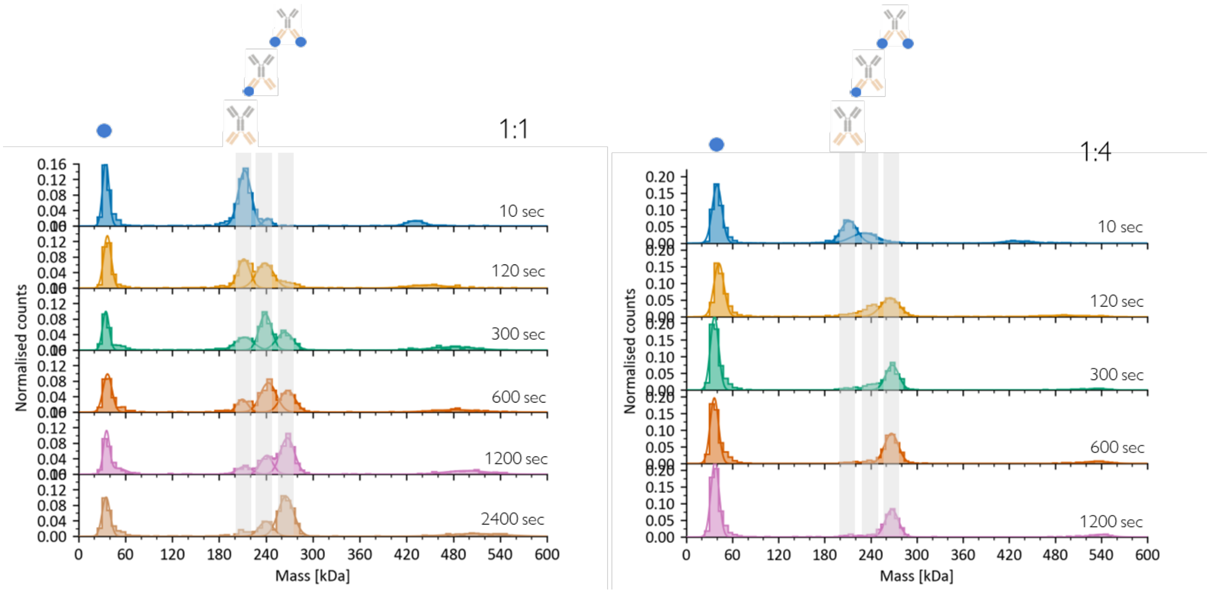


*Fig. S2 Ivonescimab:PD-1 complex formation at different molar ratios of ivonescimab and PD-1.* *Ivonescimab was incubated with PD-1 at 1:1 and 1:4 molar ratio (ivonescimab concentration was 10 nM). After indicated incubation time, mixture was diluted in PBS droplet prior to measurement.*

#### Figure S3

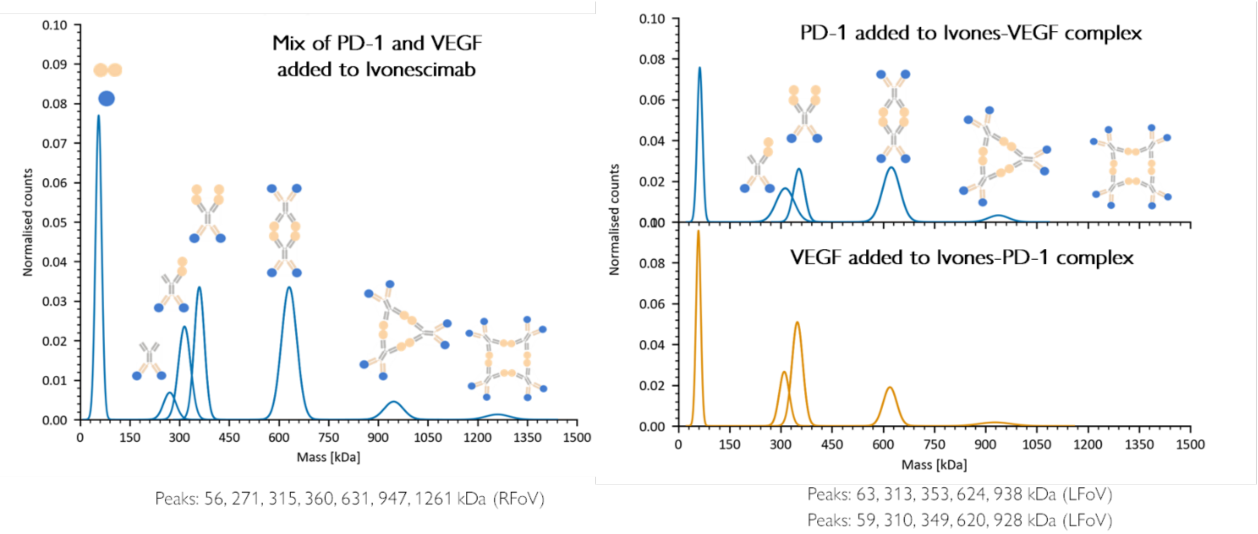


*Fig. S3 Ivonescimab:VEGF:PD-1 complex formation is stable regardless of mixing order.* *In this figure we show three different pathways of ivonescimab:VEGF:PD-1 complex formation. (A) PD-1 and VEGF mixed first. (B) Ivonescimab and VEGF mixed first. (C) Ivonescimab and PD-1 mixed first. The resulting complex profile was almost identical for all three methods. The molar ratio for all 3 reactions was 1:2:2 (50:100:100 nM). The mixtures were diluted in a PBS droplet prior to measurement.*

### Table S1

| Assumed PD-1 concentration (nM) | $K_D(1)$ (nM) | $K_D(2)$ (nM) | $k_{on}(1)$ ($nM^{-1} s^{-1}$) | $k_{off}(1)$ ($s^{-1}$) | $k_{on}(2)$ ($nM^{-1} s^{-1}$) | $k_{off}(2)$ ($s^{-1}$) |
|---|---|---|---|---|---|---|
| 15 | $7.40 \times 10^{-1} \pm 2.59 \times 10^{-1}$ | $1.56 \times 10^{-1} \pm 2.56 \times 10^{-1}$ | $4.05 \times 10^{-4} \pm 1.40 \times 10^{-5}$ | $3.00 \times 10^{-4} \pm 1.12 \times 10^{-4}$ | $2.00 \times 10^{-4} \pm 8.89 \times 10^{-6}$ | $3.13 \times 10^{-5} \pm 5.24 \times 10^{-5}$ |
| 16 | $9.21 \times 10^{-1} \pm 2.87 \times 10^{-1}$ | $3.03 \times 10^{-1} \pm 2.83 \times 10^{-1}$ | $3.78 \times 10^{-4} \pm 1.31 \times 10^{-5}$ | $3.48 \times 10^{-4} \pm 1.17 \times 10^{-4}$ | $1.84 \times 10^{-4} \pm 8.13 \times 10^{-6}$ | $5.56 \times 10^{-5} \pm 5.37 \times 10^{-5}$ |
| 17 | $1.10 \times 10^{0} \pm 3.15 \times 10^{-1}$ | $4.49 \times 10^{-1} \pm 3.09 \times 10^{-1}$ | $3.54 \times 10^{-4} \pm 1.23 \times 10^{-5}$ | $3.90 \times 10^{-4} \pm 1.21 \times 10^{-4}$ | $1.69 \times 10^{-4} \pm 7.48 \times 10^{-6}$ | $7.61 \times 10^{-5} \pm 5.48 \times 10^{-5}$ |
| 18 | $1.29 \times 10^{0} \pm 3.42 \times 10^{-1}$ | $5.95 \times 10^{-1} \pm 3.36 \times 10^{-1}$ | $3.33 \times 10^{-4} \pm 1.16 \times 10^{-5}$ | $4.28 \times 10^{-4} \pm 1.24 \times 10^{-4}$ | $1.57 \times 10^{-4} \pm 6.93 \times 10^{-6}$ | $9.36 \times 10^{-5} \pm 5.58 \times 10^{-5}$ |
| 19 | $1.47 \times 10^{0} \pm 3.70 \times 10^{-1}$ | $7.41 \times 10^{-1} \pm 3.62 \times 10^{-1}$ | $3.15 \times 10^{-4} \pm 1.10 \times 10^{-5}$ | $4.63 \times 10^{-4} \pm 1.28 \times 10^{-4}$ | $1.47 \times 10^{-4} \pm 6.46 \times 10^{-6}$ | $1.09 \times 10^{-4} \pm 5.66 \times 10^{-5}$ |
| 20 | $1.66 \times 10^{0} \pm 3.97 \times 10^{-1}$ | $8.87 \times 10^{-1} \pm 3.89 \times 10^{-1}$ | $2.98 \times 10^{-4} \pm 1.05 \times 10^{-5}$ | $4.94 \times 10^{-4} \pm 1.31 \times 10^{-4}$ | $1.37 \times 10^{-4} \pm 6.05 \times 10^{-6}$ | $1.22 \times 10^{-4} \pm 5.73 \times 10^{-5}$ |

***Table. S1 Sensitivity of $K_D(1)$ and $K_D(2)$ to assumed total PD-1 concentration in the kinetic model for the ivonescimab-PD-1 interaction.*** *For the model shown in Fig. 5, for different fixed values of the initial PD-1 concentration (column 1), the model was optimized against the empirical data, yielding the corresponding optimal values for the fitted kinetic parameters ($k_{on}(1)$, $k_{off}(1)$, and $k_{on}(2)$, $k_{off}(2)$ – columns 4-7) and the resulting $K_D(1)$ and $K_D(2)$ (columns 2-3).*
